# Young guard cells function dynamically despite low mechanical anisotropy but gain efficiency during stomatal maturation in *Arabidopsis thaliana*

**DOI:** 10.1101/2024.02.15.580511

**Authors:** Leila Jaafar, Yintong Chen, Sedighe Keynia, Joseph A. Turner, Charles T. Anderson

## Abstract

Stomata are pores at the leaf surface that enable gas exchange and transpiration. The signaling pathways that regulate the differentiation of stomatal guard cells and the mechanisms of stomatal pore formation have been characterized in *Arabidopsis thaliana*. However, the process by which stomatal complexes develop after pore formation into fully mature complexes is poorly understood. We tracked the morphogenesis of young stomatal complexes over time to establish characteristic geometric milestones along the path of stomatal maturation. Using 3D-nanoindentation coupled with finite element modeling of young and mature stomata, we found that despite having thicker cell walls than young guard cells, mature guard cells are more energy efficient with respect to stomatal opening, potentially attributable to the increased mechanical anisotropy of their cell walls and smaller changes in turgor pressure between the closed and open states. Comparing geometric changes in young and mature guard cells of wild-type and cellulose-deficient plants revealed that although cellulose is required for normal stomatal maturation, mechanical anisotropy appears to be achieved by the collective influence of cellulose and additional wall components. Together, these data elucidate the dynamic geometric and biomechanical mechanisms underlying the development process of stomatal maturation.

**Significance:** We examined how young stomata develop into mature complexes and the functional consequences of stomatal maturation, establishing geometric milestones and detecting mechanical changes in wall anisotropy and turgor pressure during stomatal maturation. We show that mature guard cells exert less energy and require a lower increase in turgor pressure to open the pore, likely due to higher wall anisotropy that results from the patterned deposition and/or remodeling of cellulose and other wall components.

## Introduction

The stomate is a dynamic system at the leaf surface that regulates gas exchange and water relations. Stomatal complexes at their simplest are composed of two guard cells separated at the center of their common boundary by a pore. Guard cells reversibly deform to regulate the size of the stomatal pore in response to environmental stimuli such as changes in light, temperature, humidity, water content, and CO_2_ levels. The responses of guard cells to opening- and closure-inducing conditions have been extensively studied in *Arabidopsis thaliana* (Arabidopsis) (Engineer et al. 2016; Kinoshita et al. 2001, 2003). The cell walls of Arabidopsis stomata contain circumferentially organized cellulose, which limits the radial expansion of these cells, causing them to elongate in response to opening stimuli (Rui and Anderson 2016; Yi, Chen, and Anderson 2022). Under opening conditions, yielding of the guard cell wall is accompanied by an increase in turgor pressure as water enters the cell (Yi et al. 2022). These combined processes cause the guard cells, which are longitudinally constrained by polar stiffening of the stomatal complex (Carter et al. 2017), to elongate and bend, leading to widening of the stomatal complex and pore opening. In contrast, under closure conditions, guard cells deflate and their cell walls contract, leading to cell shortening and straightening and closure of the pore (DeMichele and Sharpe 1973; Franks and Farquhar 2007; Rui and Anderson 2016).

Stomatal development can be divided into three phases: differentiation from epidermal stem cells, which terminates with the division of a guard mother cell into two nascent guard cells; pore formation, in which the recently divided sister guard cells partially separate at the midpoint of their common boundary to generate a stomatal pore; and finally maturation in which both the pore and the guard cells enlarge and change shape to establish the final structure of the stomatal complex. The first phase has been characterized in detail in Arabidopsis (Guo, Wang, and Dong 2021; Lopez-Anido et al. 2021). Protodermal cells first undergo a series of division and differentiation events that give rise to meristemoid mother cells. Under the influence of the transcription factor SPEECHLESS (SPCH), a meristemoid mother cell divides asymmetrically to give rise to a meristemoid cell and a stomatal lineage ground cell. The MUTE transcription factor then drives the conversion of the meristemoid cell into a guard mother cell that divides symmetrically into two sister guard cells under the influence of a third transcription factor, FAMA (Guo et al. 2021). At this stage, the guard cells share a cell wall that connects them at the middle.

In the second phase of stomatal development, sister guard cells partially separate to form a pore that facilitates gas exchange and water transpiration. During pore formation, pectin degradation occurs at the initiation site of the pore (Rui et al. 2019). Then, the pore enlarges, which requires an increase in turgor pressure. This second phase is followed by stomatal maturation in which guard cells and the pore grow to reach their final size and, hypothetically, their optimal functional state. One challenge for stomatal complexes is that they must continuously function while maturing to enable gas exchange and photosynthesis. However, the cell biological and biomechanical processes underlying stomatal development after pore formation are not well understood.

Stomatal mechanics are governed in part by the cell wall that surrounds each guard cell, but evidence suggests that cell wall synthesis continues after stomatal pore formation (Fujita and Wasteneys 2014; Rui and Anderson 2016). Atomic force microscope (AFM) measurements have demonstrated that guard cells in *Arabidopsis thaliana* become stiffer as they mature, with the poles becoming stiffer than the rest of the complex (Carter et al. 2017). The cell walls of guard cells are mechanically anisotropic (Keynia et al. 2023), which helps explain their anisotropic deformations in response to opening and closure stimuli. However, how the wall thickness and 3D biomechanics of guard cells change over the course of stomatal maturation is unknown.

Here, we used time-lapse live-cell microscopy to establish morphogenetic milestones for stomatal maturation in the cotyledons of *Arabidopsis thaliana* seedlings, finding that guard cells alter their growth patterns from dorsoventrally symmetrical elongation to establish the final pore size to asymmetric elongation to reach the final complex dimensions. A series of 3D nanoindentation experiments showed that unlike young guard cells, mature guard cells display mechanical wall anisotropy, being relatively stiffer in the circumferential than longitudinal direction. Additionally, turgor pressure is lower in mature guard cells than in young ones in both closed and open states and changes less between these states. Our model predicts that mature stomatal complexes open more efficiently than young ones. Functional analysis of young and mature cellulose-deficient stomata showed that even though cellulose is required for normal stomatal function, its patterned deposition might not be the only driver that establishes mechanical wall anisotropy in mature guard cells. Together, these results expand our understanding of how stomatal biomechanics are established for the efficient regulation of photosynthesis and water transport in plants.

## Results

### Stomata mature through a series of morphogenetic milestones, with the pore ceasing growth before the complex

To map the maturation of stomata, we selected individual stomatal complexes that had initiated pore formation in cotyledons of 4-day-old Col-0 seedlings expressing the plasma membrane marker LTI6b-GFP and followed their growth over the next five days using confocal microscopy (Fig. 1A). Stomatal complex length (b) (Fig. 1B), pore length (a) (Fig. 1C), junction length (b-a) (Fig. 1D) and the ratio of pore length-to-junction length (a/(b-a)) (Fig. 1A, E, S1) were determined over the course of four days of maturation. As time progressed, the geometric features of stomatal complexes (complex length, pore length, and pore-to-junction length ratio) increased until they reached a plateau. A one-phase association was applied to the plots of each feature as a function of seedling age to determine the timeline of the geometric milestones for stomatal maturation (Fig. 1F). The results showed that stomatal maturation is marked by four milestones; after pore formation (I), the pore enlarges as the whole complex increases in length until the pore-to-junction ratio becomes fixed around 1.03 (II), then after some continued elongation of both the pore and the complex, the pore stops elongating (III). Finally, the entire complex stops growing (IV) at a length of around 32 μm (Fig. 1F). This model is based on regression plots of complex and pore lengths to the pore-to-junction ratio for 154 stomatal complexes (Fig. 1G-I). To differentiate between young and mature stomata in later experiments, we established a threshold of a pore-to-junction length ratio of 1 based on the association analysis results (Fig. 1F-I). Even though growth is a continuum, we divided developing stomata into two groups, young and mature to test their corresponding behavioral and mechanical properties for experimental simplicity. A stomatal complex with a pore:junction ratio <1 is denoted young, whereas a stomatal complex with a pore:junction ratio >1 is denoted mature.

**Figure 1.**
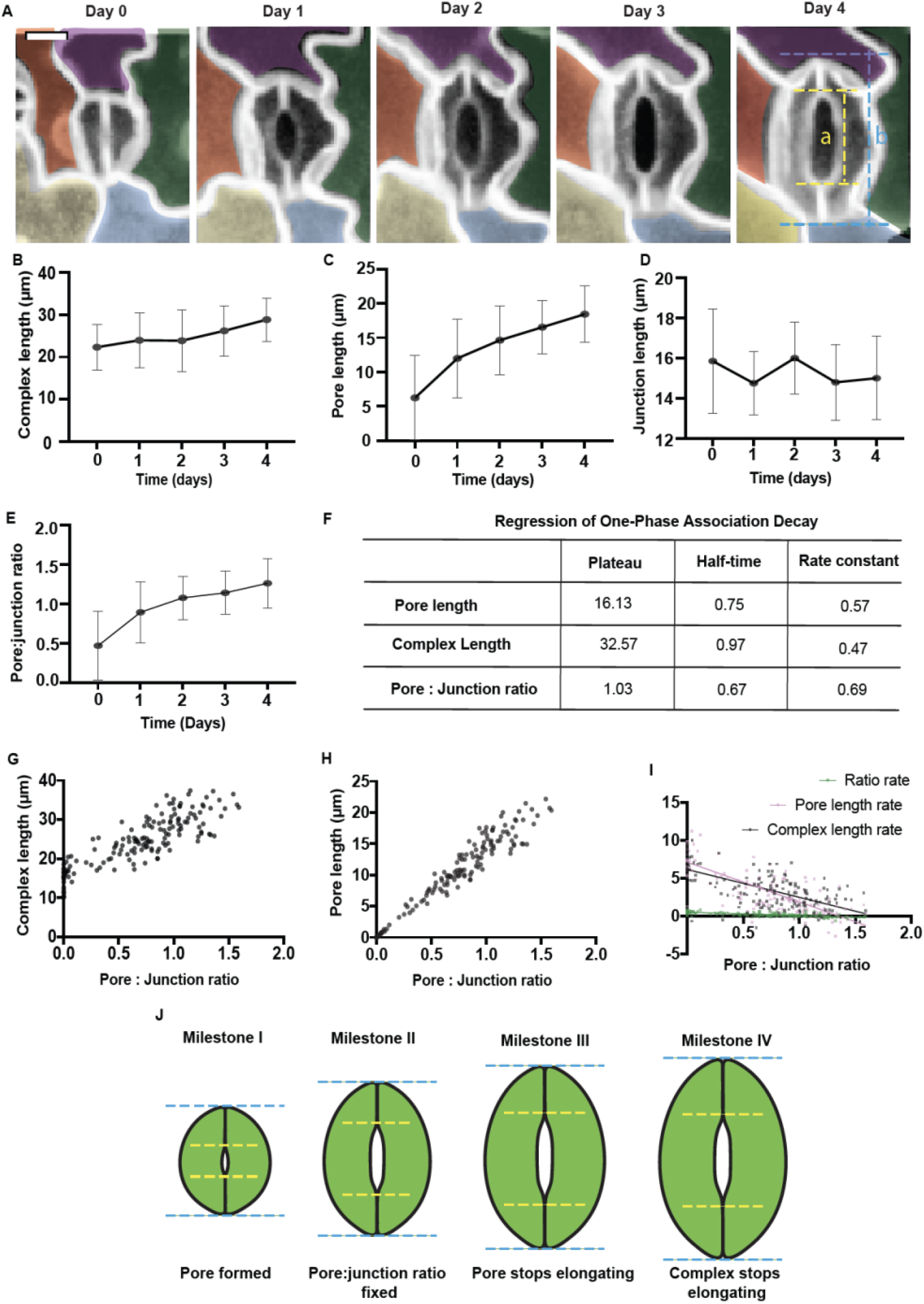
Stomata mature through a series of morphogenetic milestones. (A) Representative images of the same stomatal complex were imaged every day in Col-0 LTI6b-GFP seedlings from day four till day eight after sowing. Bar = 25 μm. (B-E) Plots of complex length (B), pore length (C), Junction length (D), and Pore: junction length ratio (a/(b-a)) (E) to the days after sowing using data collected from stomata that were done forming their pores in 4-day-old seedlings of Arabidopsis thaliana. N=5 stomata from three seedlings. Bars represent SD. (F) Regression of one-phase association decay results based on (B-E). (G-H) Plot of complex length (G) and pore length (H) to junction ratio. n = 154 stomata from six seedlings. (I) Plot of ratio rate = ((ratio on day n)-(ratio on day n-1))/1 day, pore length rate= ((pore length on day n)-(pore length on dayn-1))/1 day, and complex length rate = ((complex length on day n)-(complex length on day n-1))/1 day using data collected from stomata at all developmental stages. n = 154 stomata from six seedlings. (J) Illustration describing the identified morphogenetic milestones of stomata maturation.

### Guard cell walls thicken and gain mechanical anisotropy and turgor pressure declines during maturation

Thickening and stiffening of the cell wall at the poles of guard cells have been reported to fix the length of the stomatal complex while guard cells elongate or contract during stomatal opening and closure, respectively(Carter et al. 2017). We hypothesized that during cell maturation, other regions of the guard cell wall might thicken as well. To test whether the cell wall thickens as stomata mature, we measured wall thickness from Calcofluor white-stained confocal images of young and mature guard cells in Col-0 cotyledons (Fig. S2A). Cross-sectional views of young and mature stomatal complexes revealed overall thicker walls in mature guard cells, with a significant increase in the thickness of the inner periclinal wall (Fig. S2).

We have developed 3D nanoindentation as a tool to study the mechanical properties of wild type and mutant guard cell walls in Arabidopsis (Chen et al. 2021; Keynia et al. 2023). To analyze the mechanical properties of guard cells, nanoindentation can be performed to measure guard cell stiffness in normal, longitudinal, and circumferential directions at shallow and deep indentation depths (Fig. S3). Then these measurements can be combined with image-based measurements of individual complex geometry to build computational finite element (FE) models to determine wall modulus in different directions and turgor pressure for each measured guard cell that match the experimental nanoindentation measurments (Keynia et al. 2023). The experimental longitudinal (Fig. 2A) and circumferential (Fig. 2B) cell stiffness at shallow (300 nm) and deep (1250 nm) indentation depths show differences in young and mature guard cells (n = ten guard cells for each maturity level). Several mechanical factors contribute to measured cell stiffness: cell wall properties, cell geometry, boundary properties, and turgor pressure. To determine the wall properties and turgor pressure change that matches the experimental data, FE models of young and mature guard cells were used (Fig. 2C).

**Figure 2.**
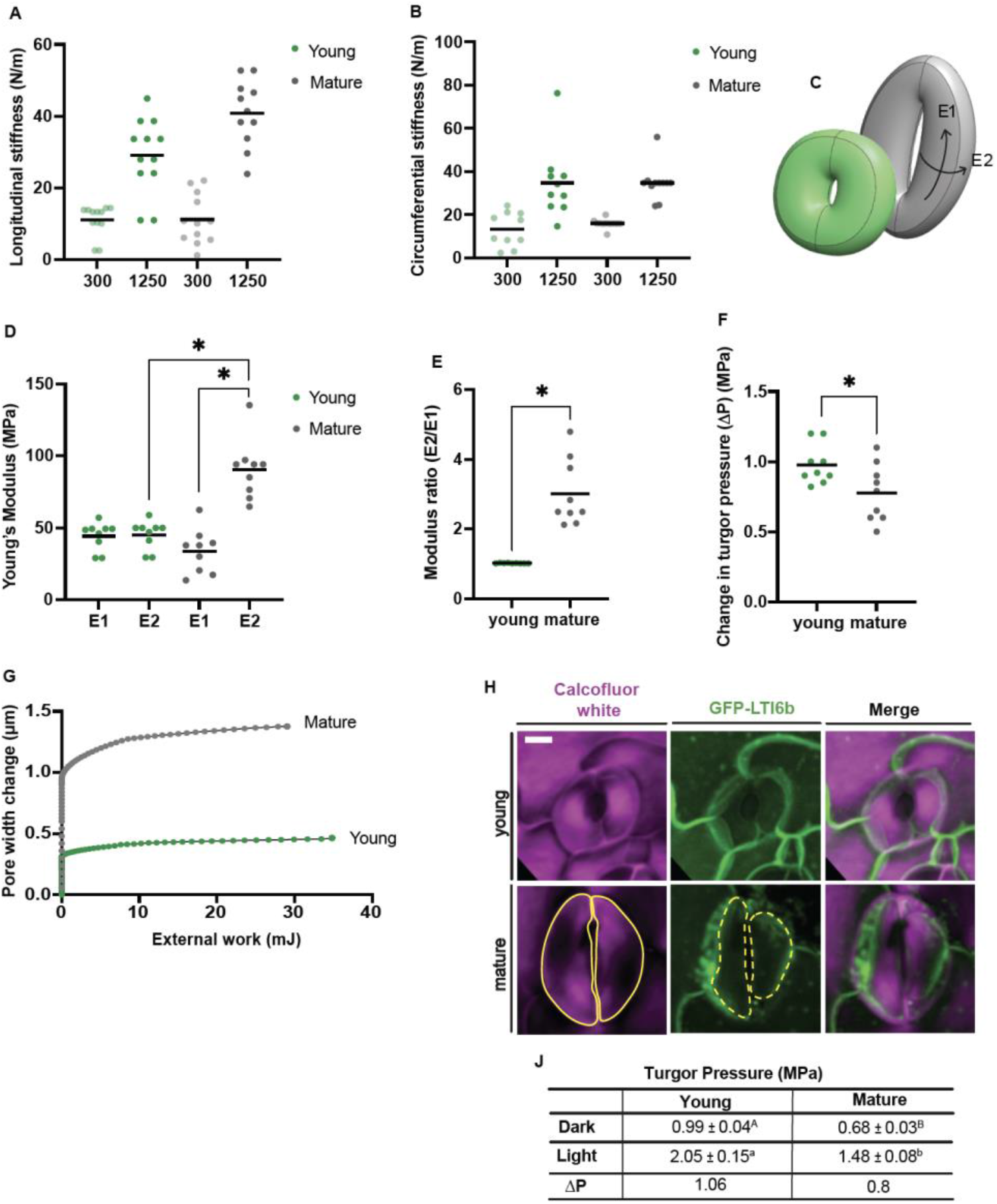
Mature guard cells display wall mechanical anisotropy and lower change in turgor pressure compared to young complexes. (A-B) Experimental apparent wall stiffness of young and mature guard cells in the longitudinal (A) and circumferential (B) directions for shallow (300 nm) and deep (1250 nm) indentation depths. n = 10 young and mature guard cells. (C) Model of young and mature stomata defining the longitudinal (E1) and circumferential (E2) modulus directions. (D) FEM estimations of young modulus of young and mature guard cell walls in longitudinal (E1) and circumferential (E2) directions. (E) Calculated young and mature guard cell wall modulus ratios (E2:E1). (F) FEM estimations of change in turgor pressure (ΔP) in young and mature stomata from closed to open states. (G) FEM predictions of the change in pore width (μm) young (green) and mature (gray) stomata as a function of the exerted external work (mJ). (H) Representative images of incipient plasmolysis outcome of young (upper) and mature (lower) stomata. Magenta and green panels correspond to the cell wall (Calcofluor white) and plasma membrane (GFP-LTI6B). Yellow lines correspond to the boundaries of the cell wall (solid yellow) and cell membrane (dashed yellow) which point to locations of membrane retraction from the cell wall (plasmolysis). Bar = 20 μm. (J) Turgor pressure estimations from incipient plasmolysis assay results of young and mature guard cells in dark (closed) and light (open) conditions. Values are presented as mean ± SE. n = 64 stomata from at least 12 seedlings. Upper (P < 0.005) and lower (P < 0.03) case letters present significance obtained from two Student’s t-tests. * P < 0.05, ANOVA.

Comparing wall modulus in the E1 (longitudinal) and E2 (circumferential) directions in young and mature stomata can provide insights into the mechanical changes guard cells undergo during maturation. We used FE analysis to predict the longitudinal (E1) and circumferential (E2) wall moduli of young and mature guard cells (Fig. 2C, D). We observed that as the guard cell matures, the longitudinal wall modulus (E1) remains relatively constant while the circumferential modulus (E2) roughly doubles, leading to a substantial increase in the circumferential:longitudinal modulus ratio (E2:E1) (Fig. 2D, E). Higher wall modulus in one direction than the other implies enhanced mechanical anisotropy and differential mechanical responses to changes in turgor pressure, with high circumferential modulus constraining guard cell widening during stomatal opening.

In addition to cell wall properties, 3D nanoindentation-FE approach derives turgor pressure changes (ΔP) upon transitioning from closed to open stomatal states. The results showed that mature guard cells display lower ΔP than young cells (Fig. 2F). Turgor pressure predictions were corroborated experimentally by incipient plasmolysis assays(Weber et al. 2015). Incipient plasmolysis is the state at which 50% of cells plasmolyze when subjected to a specific osmolyte concentration. Under the same osmotic solution, in open and closed stomatal states, a higher proportion of mature guard cells plasmolyzed compared to young guard cells (Fig. S4). The estimated turgor pressure in mature guard cells in both open and closed states was lower than that of young guard cells (Fig. 2G, H). Additionally, the change in turgor pressure from closed to opstates as estimated by incipient plasmolysis was lower in mature guard cells than in young guard cells (Fig. 2G); these data are in close agreement with the FE predictions.

Despite displaying different mechanical anisotropic properties, young and mature guard cells respond to opening stimuli by opening the pore (Fig. 3B). To identify possible reasons for gaining wall anisotropy by guard cells during maturation, we used our FE models to quantify the energy efficiency of pore opening for both ages. The results showed that mature stomata expend less energy than young stomata (29 mJ) to establish wider stomatal openings (1.377 μm) (Fig. 2G). These findings indicate that guard cells gain mechanical anisotropy in their cell walls as they mature, enabling them to achieve stomatal opening at lower turgor pressure, while using less energy.

**Figure 3.**
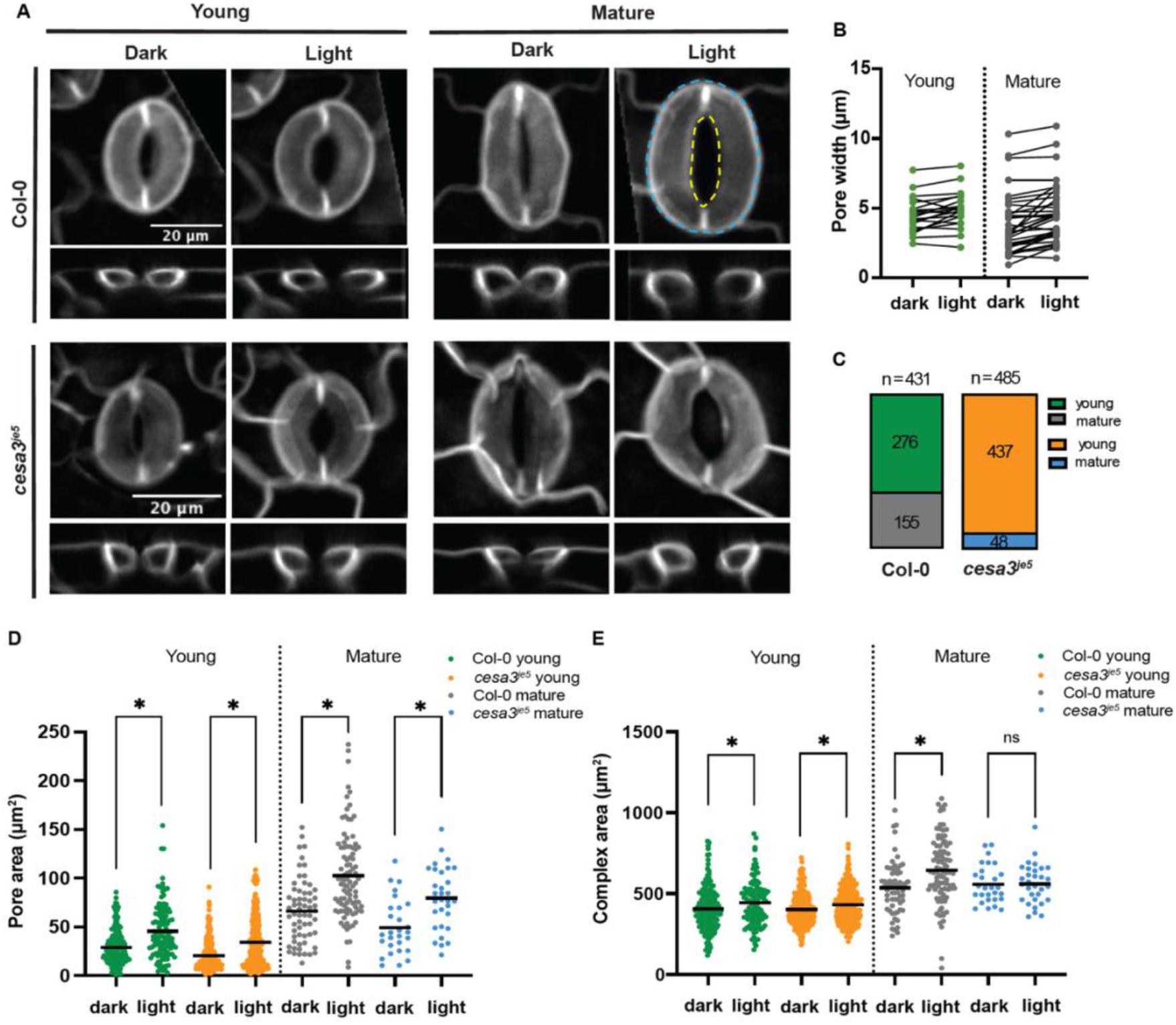
Cellulose deficient cotyledons exhibit greater percentages of young stomata with smaller relative pore size. (A) Representative images of xy (upper) and xz (lower) views of Col-0 and *cesa3*^*je5*^ young and mature stomata in response to 2.5 h of light treatment. Yellow and blue dashed lines correspond to pore and complex areas, respectively. The signal corresponds to GFP-LTI6b plasma membrane marker. Bar = 20 μm. (B) Quantification of change in pore width of young (green) and mature (gray) stomata before and after 2.5 hrs of light treatment. n > 21 stomata from eight seedlings. (C) Distribution of young (green or orange) and mature (gray or blue) stomata in Col-0 and *cesa3*^*je5*^. n = 431 stomata of Col-0 and n = 485 stomata of cesa3^je5^. (D, E) Measured pore (D) and complex (E) area of young and mature stomata of Col-0 and cesa3^je5^ in dark and light conditions. n > 62 stomata from 12 Col-0 seedlings. n > 256 young stomata and n > 27 mature stomata from 14 cesa3^je5^ seedlings. ns P > 0.05, * P < 0.05, Student’s t-test and Mann-Whitney test.

### In wild type seedlings, mature stomata open to a greater degree than young stomata

To determine the effects of higher wall anisotropy and lower turgor pressure on stomatal function, we first imaged young and mature stomatal complexes in cotyledons of Col-0 LTI6B-GFP seedlings that had been placed in the dark overnight, then subjected the seedlings to 2.5 hours of light to induce stomatal opening and imaged the same stomatal complexes, measuring geometric changes in their stomata (Fig. 3). As expected, light conditions led to stomatal opening and thus an increase in pore and complex area of both young and mature stomata (Fig. 3A-B, D-E, S6). This implies that stomatal response to opening stimuli is maintained across developmental stages. However, mature stomata displayed a greater pore:complex area ratio in dark and light conditions than young stomata (Fig. S6). This means that in Col-0, the pore occupies a larger portion of mature complexes than of young ones. Additionally, comparing the pore:complex area ratio gives insight into the degree of change in pore area with respect to complex area upon transition from closed to open stomatal states. Pore:complex area ratio did not change substantially between closed and open states in young stomata, whereas in mature stomata this ratio increased significantly (Fig. S6). Hence, although stomata at different developmental stages behave similarly in response to opening conditions, mature stomata evince a greater degree of opening, which hints at an enhanced stomatal function in regulating physiological processes such as gas exchange and water relations.

### Cellulose is required for normal stomatal maturation and achieving maximum stomatal opening

Cellulose is a major load-bearing polymer in the cell wall and has been shown to facilitate guard cell elongation in response to light (Rui and Anderson 2016). We hypothesized that the organized deposition of cellulose microfibrils might be a driver of the mechanical anisotropy in the guard cell wall predicted by nanoindentation and modeling (Fig. 2). To test this hypothesis, we determine the responses of young and mature stomata of the cellulose-deficient mutant *cesa3*^*je5*^ to light. Similar to wild-type, and despite lower cellulose content in *cesa3*^*je5*^, young and mature guard cells responded to light by opening their pores (Fig. 3A, D, S6). Interestingly, unlike wild-type controls, mature stomatal complexes of *cesa3*^*je5*^ seedlings did not show a significant change in total area when going from dark to light conditions even though their pores opened (Fig. 3 D, E). Furthermore, young and mature *cesa3*^*je5*^ stomata displayed smaller pores and a lower degree of pore opening determined by lower pore:complex area ratios relative to Col-0 (Fig. 3A, D, S6). Additionally, optical cross-sections of mature *cesa3*^*je5*^ guard cells appeared less round than Col-0 controls (Fig. 3A). We also observed that the overall number of mature stomata was low in *cesa3*^*je5*^ (∼10%), unlike Col-0 cotyledons where mature stomata presented ∼36% of the total number of stomata captured (Fig. 3C). Unlike previous reports of cellulose in the innermost layer of young guard cells being disorganized (Fujita and Wasteneys 2014), Pontamine Fast Scarlet 4B (S4B) staining, which reveals cellulose organization throughout the thickness of the wall (Anderson et al. 2010), showed that cellulose microfibrils in young and mature guard cells of Col-0 were wrapped around the cells in a radial pattern (Fig. S5). Together these data suggest that cellulose is required for normal stomatal maturation, both in terms of cell geometry and mechanical anisotropy in the wall, but that other wall components likely contribute to the functional architecture of the mature guard cell wall and can partially compensate for defects in cellulose content.

## Discussion

In *Arabidopsis thaliana*, the symmetric division of the Guard Mother Cell into two guard cells is driven by transcription factors (FAMA and FOUR LIPS) (Guo et al. 2021). After cell division, the stomatal pore is formed and enlarged by pectin degradation and increased cell turgor, respectively (Rui et al. 2019). Little is known about the mechanisms underlying stomatal maturation after pore formation. Here, we identified geometric and mechanical changes that occur during guard cell maturation and their influence on stomatal function.

We started by determining the chronological order of geometric changes of the stomatal complex when transitioning from young to mature states (Fig. 1). We tracked the same complexes for four days after pore formation and assessed changes in their geometric features including complex, pore, and junction lengths (Fig. 1A-D). Then, we fitted the geometric measurements into a one-phase association model as a function of seedling age to establish a stomatal maturation timeline (Fig. 1G-I). The model revealed four morphogenetic milestones of stomatal maturation (Fig. 1J). After pore formation (Milestone I), the guard cells grow until the pore and junctions are similar in length (pore:junction length ratio ∼1.03) (Milestone II). Guard cells continue elongating symmetrically until the inner periclinal cell region stops to fix the pore length (Milestone III). Finally, guard cells expand asymmetrically where the outer periclinal cell region expands to increase the complex length, then stops when it reaches the size of a fully mature stomatal complex (Milestone IV). The samples were moved from the microscope to the growth chamber continuously for sequential observation. However, the consistent results from different seedlings overcome the variability that might be introduced by the experimental design.

Microscopy evidence shows that as guard cells mature, their cell walls thicken and become organized radially (Fujita and Wasteneys 2014; Zhao and Sack 1999), potentially driving the longitudinal expansion of guard cells in response to opening stimuli and constraining their circumferential widening. While young guard cells display similar wall thickness around the cell diameter, the inner periclinal wall is thicker than the outer periclinal wall in mature guard cells, consistent with prior measurements of wall thickness using transmission electron microscopy (Keynia et al. 2023) and we also observed thicker walls in the ventral anticlinal region of mature guard cells (Fig. S2). AFM measurements across the width of the stomatal complex have shown that the anticlinal wall facing the pore is stiffer than the dorsal anticlinal wall of mature guard cells, unlike in young complexes that showed lower, roughly equal stiffness values across both regions (Carter et al. 2017). The asymmetric guard cell expansion that occurs from Milestones III to IV, where the pore stops elongating but the guard cells continue to elongate, might be explained by the increased thickness and stiffness of the ventral anticlinal wall in mature guard cells that would resist elongation in response to increasing turgor pressure during growth (Carter et al. 2017). Hence, if a thicker ventral anticlinal wall is not necessary for guard cell opening as reported by (Carter et al. 2017), it might instead be important for establishing the mature stomatal form and optimizing the efficiency of stomatal dynamics.

Previous reports highlighted the importance of computational models to determine the mechanical properties of biological systems (Boudon et al. 2015; Forouzesh et al. 2013; Hayot et al. 2012; Li et al. 2022; Woolfenden et al. 2017). We used nanoindentation coupled with finite element (FE) modeling to characterize the mechanical traits of young and mature guard cells, including cell wall moduli in different directions and turgor pressure (Fig. 2). Normal indentation at different depths provides the stiffness properties of the material in a direction perpendicular to the surface of the cell. However, lateral indentation yields in-plane mechanical aspects of the wall in different directions, which refines our understanding of the guard cell wall as a dynamic entity and provides insight into variations in turgor pressure (Keynia et al. 2023). We also used this measurement and modeling framework to calculate the work required to open stomata of different maturation states.

Using lateral nanoindentation in the longitudinal and circumferential directions, we determined cell stiffness in young and mature stomatal complexes at different depths (Fig. 2A, B, S3). To differentiate between the mechanical factors contributing to cell stiffness, we built FE models based on the geometry of young and mature guard cells, followed by an iterative process to match the modeled pore size to experimental data. The results show that while young guard cells display similar wall moduli in all directions (E2/E1 ≈ 1), mature guard cell walls present higher circumferential than longitudinal modulus (E2/E1 > 2) (Fig. 2C-E). This finding suggests that guard cells acquire anisotropic wall properties as they mature. We speculate that wall anisotropy in mature guard cells arises from the radial arrangement of the cellulose microfibrils (Fujita and Wasteneys 2014; Rui and Anderson 2016). In addition to determining cell wall properties, we employed our approach to determine turgor pressure values (Fig. 2F). The models showed a significantly smaller increase in turgor pressure change (ΔP) upon pore opening in mature guard cells than in young guard cells (Fig. 2F). These results were comparable to estimations of turgor pressure based on incipient plasmolysis experiments (Fig. 2G, H).

Measuring pore and complex area of young and mature stomata allowed us to assess their behaviors in response to light (Fig. 3). The data show that at all developmental stages, stomata respond to light by opening their pores, which increases pore and complex areas (Fig. 3A, B, D, E). These results imply that similar dynamic behaviors are achieved by cells of different sizes and maturation states in response to an opening stimulus. These experiments were done on cotyledons growing in culture (∼100% humidity), which might underestimate stomatal response to opening or closing stimuli. However, the results show a clear difference in young and mature stomatal performance that might be even greater in true leaves. Open pores in mature stomata were larger (Fig. 3A, B, D, S6). This could be simply because the mature guard cells are larger in size. However, according to the maturation milestone model (Fig. 1J), mature guard cells have a longer inner anticlinal wall and a shorter junction (pore-to-junction length >1) and a greater pore area-to-complex area ratio (Fig. S6). Therefore, the pore occupies a greater portion of the mature stomatal complex, which would be expected to enhance gas exchange across the stomatal pore, unlike young complexes that possess smaller pores relative to their complex areas (Fig. S6). This increase in pore:complex area might be a key functional feature of stomatal maturation.

Mature guard cells undergo smaller changes in turgor pressure than young guard cells yet open fully in response to opening stimuli. This might be due to the mechanical wall anisotropy gained by mature guard cells, which might aid in achieving maximum pore opening while exerting lower energy (Fig. 2G). Previous reports observed a random organization of fibrillar material at the innermost cell wall face in guard cells prior to pore formation and a radial pattern of material in mature guard cells using field emission scanning electron microscopy (FESEM) (Fujita and Wasteneys 2014); however, microtubules and thus presumably the trajectories of cellulose synthases were observed to be radial in both nascent and mature guard cells (Rui and Anderson 2016). Hence, we suspect that cellulose is a key driver of mechanical wall anisotropy and anisotropic morphogenesis in mature guard cells. However, in the abovementioned FESEM images of the innermost cell wall layer of young and mature Col-0 stomata, cellulose is not specifically labeled and the sample preparation did not include pectin degradation (Fujita and Wasteneys 2014). Furthermore, detailed analyses of cellulose synthase particle trajectories have not been performed in young and mature guard cells. Moreover, to our knowledge, a clear morphological boundary between young and mature stomata has not been reported. Therefore, even though the abovementioned images are insightful and provide a useful reference for wall patterning at the innermost layer in different types of cells, we cannot yet make definitive conclusions regarding cellulose arrangement in young and mature guard cells.

Our results with a cellulose synthase mutant provide evidence that cellulose contributes to the mechanical anisotropy of mature guard cells but is not the sole determinant of this anisotropy. The radial arrangement of cellulose in young and mature stomata determined by S4B staining and the ability of cellulose-deficient guard cells to close and open the pores in response to dark and light conditions (Fig. 3), means that other cell wall components besides cellulose likely contribute to the mechanical wall anisotropy determined by our approach (Fig. 2). Pectin is a major component of the cell wall that contributes to guard cell function (Chen et al. 2021; Rui et al. 2017) and wall anisotropy in mature guard cells (Keynia et al. 2023). Overexpression of *POLYGALACTURONASE INVOLVED IN EXPANSION3* (*PGX3*), a pectin-degrading enzyme, affects the speed of stomatal response (Rui et al. 2017), and results in higher mechanical anisotropy in the guard cell wall, whereas knocking out *PGX3* results in lower wall anisotropy as determined by lateral nanoindentation (Keynia et al. 2023). Similar to *PGX3* knock-out mutant, cellulose-deficient mutant *cesa3*^*je5*^ displayed lower mechanical anisotropy as determined from the nanoindentation-FE approach. These results imply that mechanical wall anisotropy in mature guard cells is achieved by the combined effects of cellulose and pectin such that the pectin stiffness profile across the guard cell must have an orientation dependence. It is unclear at this time whether the pectin anisotropy is related to the cellulose through some type of cross-linking or if it is the result of anisotropy during pectin synthesis. Further mechanical studies are needed to identify additional differences between young and mature guard cells of cell wall mutants that can be correlate with altered stomatal function.

In summary, our data depict the morphogenetic and biomechanical milestones of stomatal maturation in *Arabidopsis thaliana*. After pore formation, the whole complex grows until the pore reaches its maximum length. The guard cells continue to elongate asymmetrically to establish the final mature form of the stomatal complex. Mature guard cells gain mechanical anisotropy in their walls through the organization of cellulose and other wall components. Our modeling results indicate that the acquisition of mechanical anisotropy in the guard cell wall enables an energy-efficient stomatal response to environmental stimuli (Fig. 4). Further studies can build on this biomechanical and morphogenetic foundation to dissect additional genetic and molecular factors that influence how efficient stomatal function is maintained and enhanced over the course of the final stage of stomatal development.

**Figure 4.**
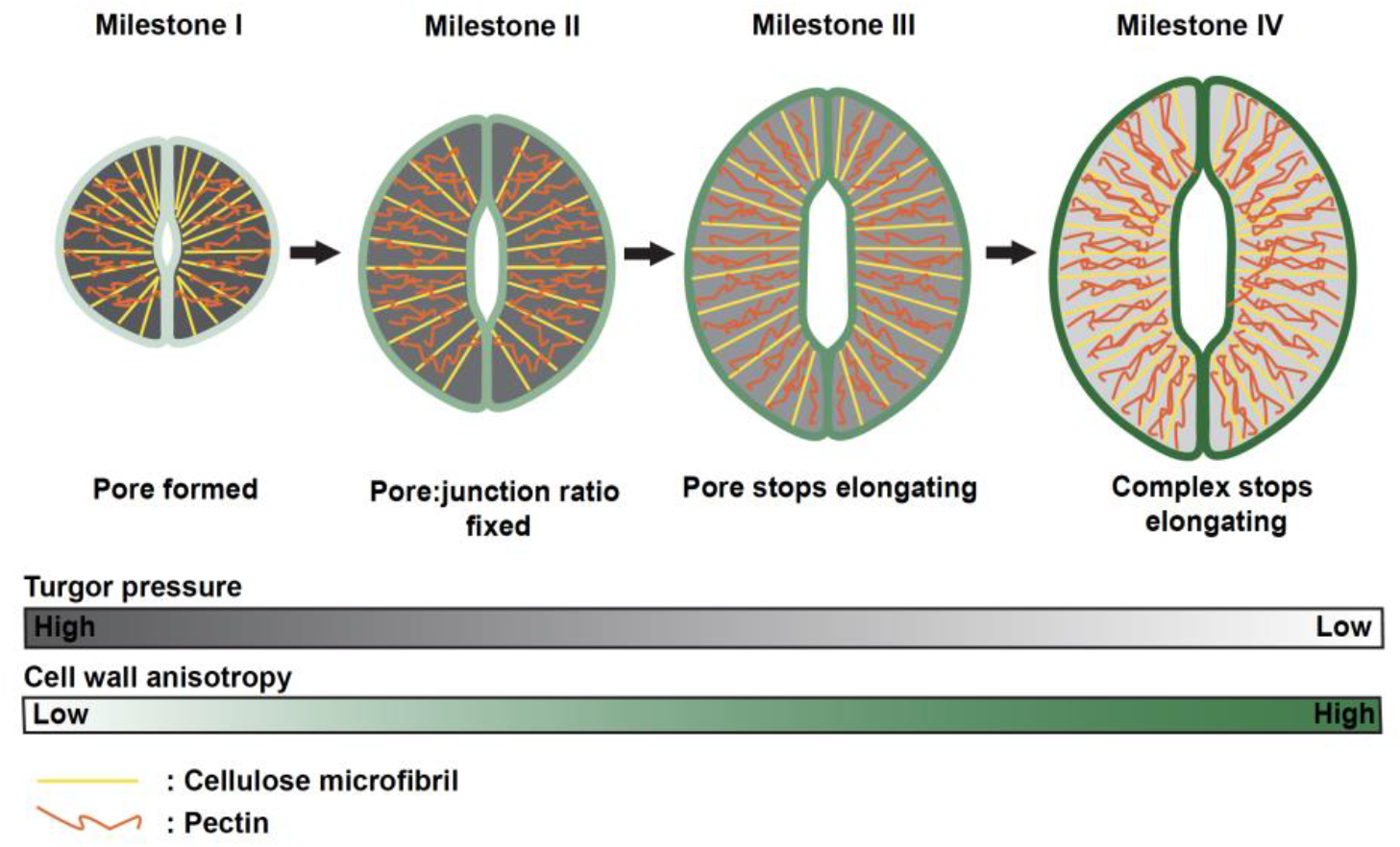
Stomata mature through a series of geometric milestones. Stomatal maturation starts with the formation of the pore that elongates along with the guard cells until pore: junction ratio gets fixed at > 1. Then the pore stops elongating while the complex grows to achieve the final mature stomatal form. During maturation, guard cell turgor pressure (gray) decrease while cell wall anisotropy (green) enhances. This is achieved by the organization of cellulose (yellow lines) and other cell wall components like pectin (orange lines).

## Materials and Methods

### Plant Growth Conditions

*Arabidopsis thaliana* Col-0 LTI6B-GFP (Cutler et al. 2000) and *cesa3*^*je5*^ LTI6B-GFP seeds were sterilized in 30% bleach and 0.1% SDS for 20 min then stratified in 0.15% (w/v) agar at 4°C in the dark for at least three days. Seeds were sowed on Murashige and Skoog (MS) plates containing 2.2 g/L MS salts (Caisson Laboratories, North Logan, UT, USA), 0.6 g/L MES, 1% (w/v) Sucrose, and 0.8% (w/v) agar (Sigma, St Louis, MO, USA), pH 5.6. Seedlings were grown at 22°C under 24 h light (4100K fluorescent Lamp, 800, 900 PPFD).

### Guard Cell Tracking

We selected stomata that had just formed pores from cotyledons of four-day-old Col-0 LTI6B-GFP seedlings (Cutler et al. 2000) and tracked their growth for four days. Every day, the same seedlings were mounted gently on slides and imaged using a Zeiss Cell Observer SD microscope with a Yokogawa CSU-X1 spinning disk head and a 40x oil objective. Intact seedlings were then returned to the MS plate and allowed to grow for another day before being mounted and imaged again. In each sample, the same region was imaged every day until day eight after sowing. Complex, pore, and junction lengths were measured using ImageJ. We used a 488 nm excitation laser and a 525/50 nm emission filter to detect the GFP signal in all experiments.

### Instrumented Indentation Testing (Nanoindentation)

Nanoindentation experiments were conducted using a Hysitron TI Premier Nanoindenter (Bruker, USA), equipped with a 50X objective to target guard cells accurately. The conical-type tip of the probe had a diameter of 2-3 μm, and a 2 μN engaging force was applied to the middle of each cell. The input load function was defined as the displacement values, with the loading and unloading and the lateral motion rate set to 30 nm/s. During lateral indentation, the normal displacement was set to zero with respect to the previous position, while for normal indentation, the lateral displacement was set to zero to ensure that the probe would indent the center of the guard cell. The experimental normal and lateral stiffness were measured using nanoindentation as detailed in (Keynia et al. 2023) and were subsequently matched in the iterative process of the finite element (FE) method in order to determine the cell wall moduli and turgor pressure change.

### Finite Element Modeling

We followed finite element (FE) modeling methods reported in (Keynia et al. 2023). FE analysis of nanoindentation measurements was conducted using commercial finite element software (Abaqus, 2019) to estimate the wall modulus and turgor pressure of the cell in young and mature guard cells. At least nine guard cells for each developmental stage were modeled with the finite element method. Using images from a laser confocal microscope, stomatal complex length, guard cell geometry, and pore width were measured, and a model of each guard cell was constructed individually using the lofting method in SolidWorks. The initial pore width used in the FE analyses was based on the pore width at the closed state for young and mature Col-0 stomata. The thickness distribution of the cross-section of the guard cells was set based on measurements from Calcofluor images, then the structural model was imported into Abaqus. To simulate the nanoindentation measurements, the conical tip was scanned using a confocal microscope and its geometry was also imported into Abaqus. A linear anisotropic elastic model using a discrete coordinate system was assigned uniformly across the whole cell and based on the orientations of cellulose and matrix polysaccharides in the guard cell wall, the anisotropic modulus was assumed to have a relation E_1_=E_3_ and its value was calculated based on iteration with an appropriate initial guess. E2 defined the wall modulus along the circumferential direction of the cell; the modulus in this direction had the maximum value based on initial assumptions (Fujita and Wasteneys 2014; Lucas, Nadeau, and Sack 2006; Rui and Anderson 2016). Poisson’s ratios were set to ν_12_ = ν_23_= 0.3 and ν_13_ = 0.47. Shear modulus was assumed to have a relation G_12_ = G_23_, and G_13_ can be determined by G_13_ = E_1_/ [2(1 + ν_13_)]. Due to lateral indentation, contact properties such as friction coefficient can play a significant role in the stiffness values of the cell wall in lateral directions (x and y) derived from the computational model. To exclude these parameters from the iteration, lateral indentations starting from 100 nm displacement from the center and increasing up to 5 μm were performed on nine cells along the longitudinal direction to investigate the contact properties between the guard cell and nanoindenter tip during lateral motion. These experiments revealed the point at which sliding begins, the maximum lateral force, and the slope of the lateral force vs normal force (friction coefficient) of non-sliding indentation. As a result, for all the computational models the friction coefficient was set to 0.3, and the shear stress limit and fractions of characteristic surface dimensions were set to 0.3, 50 MPa, and 0.9 respectively (Keynia et al. 2023). For boundary conditions, the materials at the polar positions were confined, ventral edges were free of constraint, and dorsal edges were constrained in the vertical direction to represent constraints from adjacent pavement cells (Chen et al. 2021).The analysis was conducted in three steps: cell pressurization, normal and lateral nanoindentation. The pore width at the end of the pressurization and the stiffness at shallow and deep indentation depths were used to compare with experimental measurements iteratively (Keynia et al. 2023). Turgor pressure and wall modulus were estimated based on the simulations with consistent pore width and normal and lateral stiffness. External work, in the context of hydrostatic pressure causing cell expansion, refers to mechanical work done by the cell wall as it pushes against an external force (e.g., turgor pressure in neighboring cells) during its expansion and cell opening. The external work was an output from the FE model. The change in turgor pressure for these simulations occurred for a duration that was ten times the wall viscoelastic time constant assumed for the material model.

### Incipient Plasmolysis Assays

We followed the incipient plasmolysis assay protocol reported by Weber and Colleagues (Weber et al. 2015). Briefly, cotyledons were excised from 8-day-old Col-0 LTI6b-GFP seedlings (Cutler et al. 2000) and subjected to light or dark conditions to generate open and closed stomata sample groups, respectively. Cotyledons with open or closed stomata were soaked in 2% (w/v) Calcofluor white (Sigma, St Louis, MO, USA) in sorbitol solutions of different concentrations (0.4 M, 0.6 M, and 0.8 M for the open group, and 0.2 M, 0.3 M, and 0.4 M for the closed group) for 40 min before imaging using a Zeiss Cell Observer SD microscope with a 63X oil objective. Excitation lasers at 488 nm and 405 nm with 525/50 nm and 450/50 nm emission filters were used to detect GFP and Calcofluor white signals, respectively. Quantification and calculation of the percentage of plasmolyzed young and mature stomata were done using ImageJ. The percentage of plasmolyzed cells was plotted as a function of sorbitol concentration to perform linear regression analysis. The sorbitol concentration of incipient plasmolysis was determined (50% of guard cells are plasmolyzed) and used to calculate turgor pressure according to Van’t Hoff equation Ψ =c•R•T, where c is the concentration of sorbitol, R is the ideal gas constant (8.314 kPa•L/mol•K), and T is the temperature in Kelvin (298 K). Data were collected from nine seedlings in total in three independent experiments per condition.

### Stomatal Dynamics Assays

To compare the response of young and mature stomata to light, ten-day-old Col-0 LTI6B-GFP seedlings (Cutler et al. 2000) were incubated in the dark for 24 h to induce stomatal closure. Dark-treated cotyledons were excised and Zeiss Cell Observer SD microscope with a Yokogawa CSU-X1 spinning disk head and a 40x oil objective was used to image stomatal complexes. Then, the cotyledons were gently removed from the slide and incubated in stomatal opening solution (50 mM KCl, 0.1 mM CaCl2, and 10 mM MES-KOH, pH 6.15) and light for 2.5 h to induce stomatal opening. The same cotyledons were mounted on slides and the same regions were imaged again to capture stomata in open states. ImageJ was used to measure changes in pore width before and after light treatment. Eight seedlings were imaged in one experiment.

For a population-level assessment of young and mature stomatal function, eight-day-old Col-0 LTI6B-GFP or *cesa3*^*je5*^ *LTI6B-GFP* seedlings subjected to dark overnight then moved to light conditions for 2.5 h to induce stomatal closing and opening, respectively. Excised cotyledons were imaged using Zeiss Cell Observer SD microscope with a Yokogawa CSU-X1 spinning disk head and a 40x oil objective with 488 nm excitation and a 525/50 nm emission filter. ImageJ was used to evaluate pore and complex area of closed or open young and mature stomata. Nine to 12 seedlings were imaged in three independent experiments.

### Pontamine Fast Scarlet 4B (S4B) Staining

Eight-day-old Col-0 seedlings were grown in continuous light till day seven where they were incubated in the dark overnight to induce stomatal closure. Then seedlings were moved to light conditions for 2.5 h to induce stomatal opening. Cotyledons from seedlings in dark and light conditions were excised and cleared to disrupt the cuticle following the tissue-clearing protocol in (Sharma 2017). Briefly, cotyledons were incubated in clearing solution (7:1 95% ethanol:acetic acid) overnight at room temperature. Then, the clearing solution was removed, and cotyledons were incubated in 1M KOH for 30 min to 1 h. Cotyledons were then washed twice with water and stained with 0.1% (w/v) S4B (Sigma, St Louis, MO, USA)(Anderson et al. 2010) in liquid MS medium for 30 min at room temperature. Finally, S4B-stained cotyledons were mounted on slides for confocal imaging. Z-stacks were collected using a Zeiss Cell Observer SD microscope with a Yokogawa CSU-X1 spinning disk head and a 100x oil objective and a 561 nm excitation filter. Confocal z-series were recorded with a 0.2 μm step size. Raw confocal z-stacks were deconvoluted in AutoQuant X2 (Media Cybernetics) and maximum projections of the deconvoluted images were obtained using ImageJ.

## Cell Wall Thickness Measurements

Eight-day-old Col-0 LTI6b-GFP seedlings (Cutler et al. 2000) were stained with 2% (w/v) Calcofluor white (Sigma, St Louis, MO, USA) for 5 min at room temperature. Z-stack images of young and mature stomata were collected using Zeiss Cell Observer SD microscope with a Yokogawa CSU-X1 spinning disk head and a 63x oil objective. Z-stacks were deconvoluted using AutoQuant X2 and the thickest regions of the outer and inner periclinal walls were measured using ImageJ. Nine seedlings were imaged in three independent experiments.

## Statistical Analyses

Statistical analysis was done using Graphpad. Dots represent individual data points and lines represent mean value of the set. Sample size, statistical significance annotation (asterisks), and statistical tests corresponding to each plot are described in figure legends.

## Acknowledgments

Thanks to Hojae Yi, James Z. Wang, and Dolzodmaa Davaasuren for helpful discussions. This work was supported by collaborative U.S. National Science Foundation grant MCB-2015943 (CTA)/MCB-2015947 (JAT).

