## Supplemental figures for "Young guard cells function dynamically despite low mechanical anisotropy but gain efficiency during stomatal maturation in *Arabidopsis thaliana*"

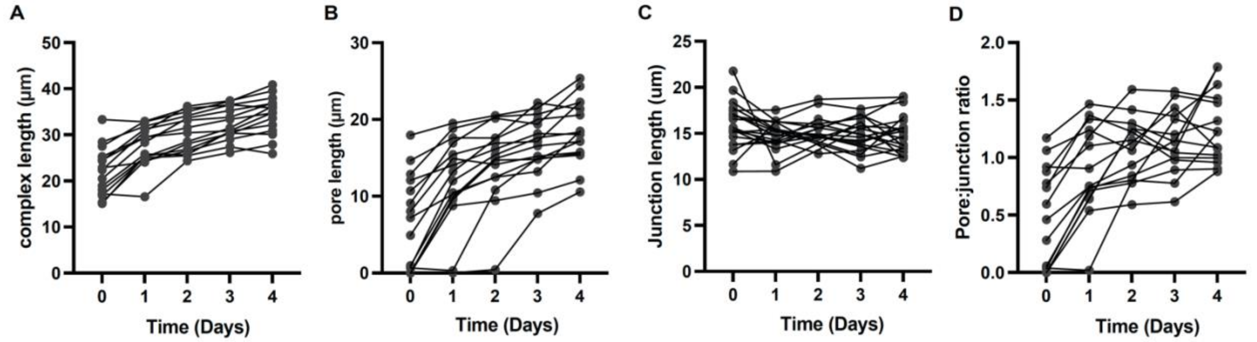


**Figure S1. Stomatal maturation trends in Col-0 cotyledons.** Tracking stomatal maturation by measuring complex (A), pore (B), and junction lengths (C) over time (days). (D) calculated pore-to-junction ratio over time (days). n = 15 stomata from three seedlings.

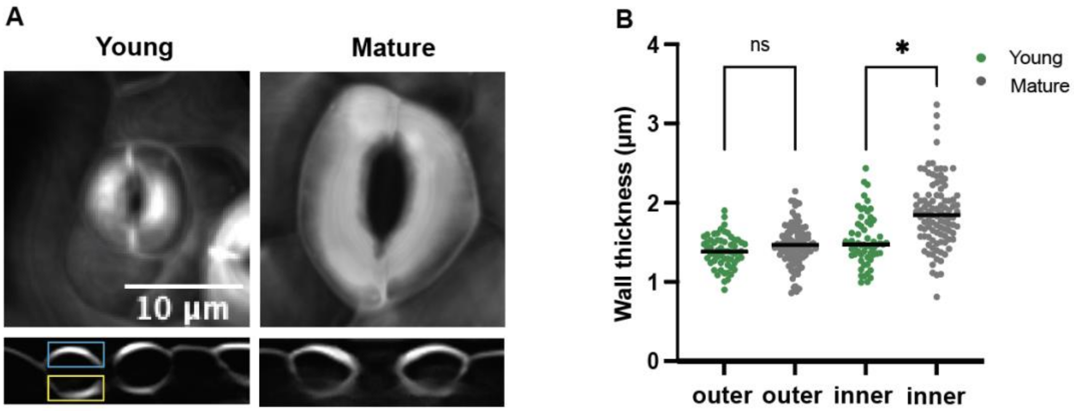


**Figure S2. Mature guard cells have a thicker inner periclinal cell wall.** (A) Calcofluor white staining of young and mature guard cells in xy (upper) and xz (lower) projections. Yellow and blue boxes highlight the inner and outer periclinal walls, respectively. Bar = 10 µm. (B) Quantification of inner and outer periclinal wall thickness in young (green) and mature (gray) guard cells. n > 60 stomata from nine seedlings. ns > 0.05, * P <0.05, Student's t-test.

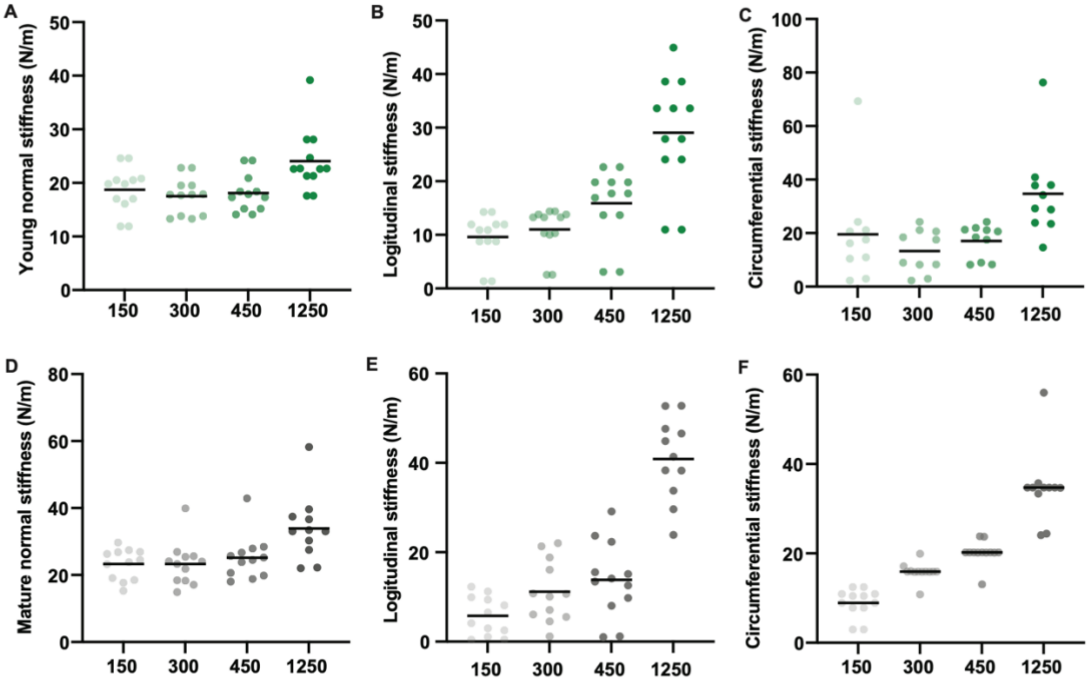


**Figure S3. Measurements of stiffness of young and mature guard cells using nanoindentation at different depths.** (A-C) Apparent stiffness measurements of young guard cells in normal (A), longitudinal (B), and circumferential (C) directions using nanoindentation at various indentation depths (150 nm, 300 nm, 450 nm, and 1250 nm). n > 10 guard cells in each category. (D-F) Apparent stiffness measurements of mature guard cells in normal (A), longitudinal (B), and circumferential (C) directions using nanoindentation at various indentation depths (150 nm, 300 nm, 450 nm, and 1250 nm). n > 10 cells per category.

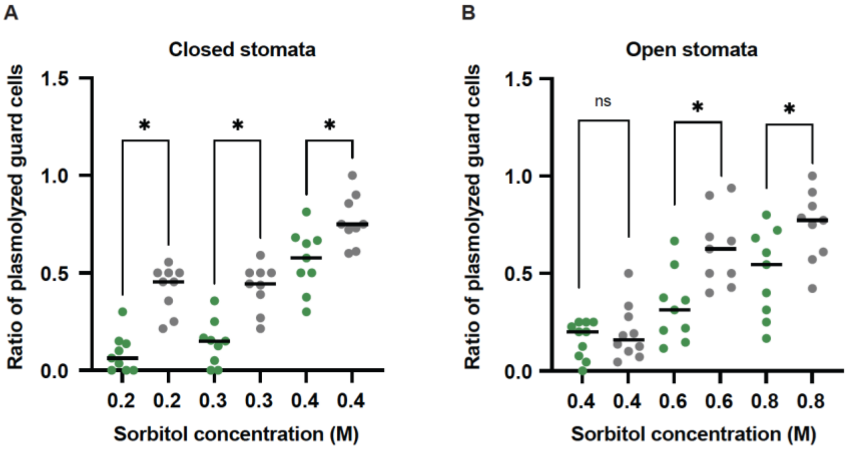


**Figure S4. Ratios of plasmolyzed young and mature guard cells in closed and open states as a function of sorbitol concentration.** (A-B) Quantification of the ratio of plasmolyzed young (A) and mature (B) guard cells at different osmotic concentrations (M) in dark (closed) and light (open) conditions. n > 64 guard cells from at least nine seedlings. ns P > 0.03, * P < 0.03, Student's t-test.

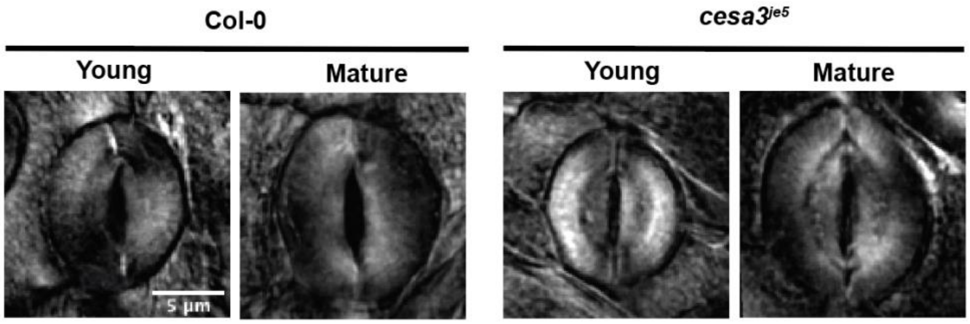


**Figure S5. Cellulose microfibrils are circumferentially wrapped around young and mature guard cells of Col-0 and *cesa3*$^{je5}$ stomata.** Maximum projections of deconvoluted z-series of S4B-stained young and mature guard cells of Col-0 (left) and *cesa3*$^{je5}$ (right) seedlings. Bar = 5 µm.

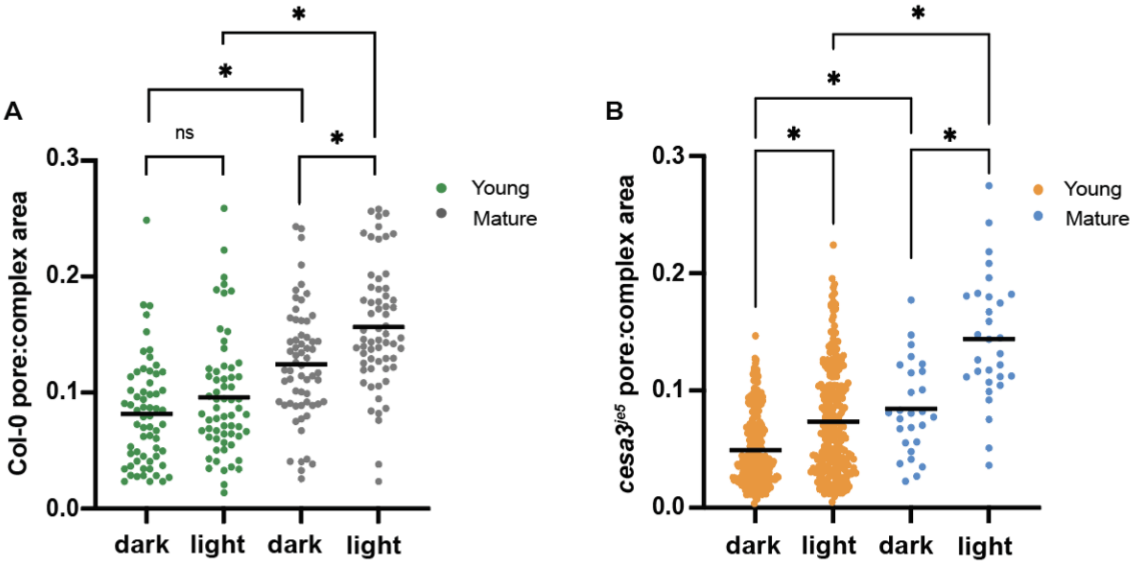


**Figure S6. Mature stomata open to a greater degree than young stomata in Col-0 seedlings, and relative pore sizes in young and mature stomata of *cesa3*$^{je5}$ seedlings (related to Figure 3).** (A, B) pore: complex area ratios of young (green or orange) and mature (gray or blue) stomata of Col-0 (A) and *cesa3*$^{je5}$ (B) seedlings before and after 2.5 h of light treatment. n = 62 young and mature stomata from 12 Col-0 seedlings. n > 255 young stomata and n > 28 mature stomata from 14 *cesa3*$^{je5}$ seedlings. ns P > 0.05, * P < 0.05, Student's t-test and Mann-Whitney test.
